# Mutation ages and population origins inferred from genomes in structured populations

**DOI:** 10.1101/2025.01.28.635333

**Authors:** Anna A. Nagel, Bruce Rannala

## Abstract

Inferring the time of origin (age) of mutations is an old question in population genetics and inferring their population of origin has become of particular interest with the sequencing of the Neanderthal genome. However, existing methods to infer mutation ages and populations of origin do not explicitly consider population structure, migration rates, and divergence times, which may bias estimates, and it is unclear how to even apply single-population estimators to structured populations. We develop a method to jointly estimate the time and population of origin of a mutation (as well as the ancestral and derived states) in a structured population using population genomic data and examine its statistical performance using simulations. Results indicate that mutation age and population of origin can be quite uncertain, even with long sequences or many samples, but this uncertainty is accurately captured using credible intervals/sets. The ancestral nucleotide state is relatively easy to infer. We apply our method to whole genome data from the 1000 Genomes Project, analyzing seven SNP mutations from six genes associated with human skin pigmentation for populations from Great Britain, China, and Kenya. Our results partially support previous conclusions, with the putative ancestral alleles from the literature matching our inferences, while the mutation age estimates only overlap in some cases. Furthermore, there are non-trivial posterior probabilities of recurrence for three of the mutations.

## Introduction

Population geneticists have long studied historical properties of individual mutations (traditionally referred to as alleles), such as whether a particular allele is ancestral to all other alleles in a population and the time in the past at which it arose (reviewed by Slatkin and Rannala 2000). Classical population genetics approaches consider the relationship between the population or sample frequency of an allele and its age (Kimura and Ohta 1973; Griffiths and Tavaré 1998), or the probability that it is the ancestral allele (Ewens and Gillespie 1974; Watterson and Guess 1977). Allele frequency is only weakly informative about allele age, especially when mutation rates increase (Ewens and Gillespie 1974); the development of genetic markers closely linked to mutations (and subsequently complete adjacent sequences) allowed the local genealogy of a sample of sequences flanking a mutation to be used to infer age, which is potentially more informative than frequency alone (Slatkin and Rannala 2000). In principle, both recombination and mutation in adjacent sequences can be modeled to develop estimators of mutation/allele age using the ancestral recombination graph (ARG) but this is computationally intensive and approximate methods have instead been developed that are most accurate for use with rare mutations (Slatkin and Rannala 1997; Rannala and Slatkin 1998; Rannala and Reeve 2002).

Existing methods for inferring allele ages and ancestral allele status typically assume a single panmictic population. How-ever, many alleles are shared between populations that are semi-isolated; in particular, human populations have a history of populations splitting from common ancestral populations with subsequent gene flow between them. Allele age estimates that do not account for population subdivision may be biased. A question of related interest, also not currently addressed, is the population of origin for a particular non-recurrent mutation. A case of obvious interest here is introgression between Neanderthal, Denisovan, and modern humans (Green *et al*. 2010) and the question of which human alleles arise from Neanderthals. Hidden Markov model-based methods have been proposed to identify Neanderthal alleles present in individual human genomes, for example, but they do not explicitly model population structure, divergence times, and migration rates (Sankararaman *et al*. 2016; Bergström *et al*. 2020). Some studies suggest that Neanderthal alleles may be risk factors for disease, for example, severe COVID-19 (Zeberg and Pääbo 2020). It is also of interest whether recurrences of mutations have occurred in different human populations – this could suggest a selective advantage for a mutation. Methods have recently been proposed to infer the complete vector of genealogies relating human individuals across the genome (the ARG) which can also potentially identify recurrent mutations (Wohns *et al*. 2022) but the methods are computationally intensive and it is difficult to quantify the uncertainty of the estimates; there is a need for a more focused approach targeting particular mutations of interest and generating reliable measures of statistical uncertainties.

In this paper, we develop a Bayesian inference method, implemented in the software program MutAnce, that exploits outputs from the BPP (Flouri *et al*. 2018) phylogeography program – which implements a structured multispecies coalescent model – to allow estimation of ages, recurrences, and populations of origin for individual alleles at single nucleotide polymorphisms (SNPs) using samples of genome sequences of individuals. The method provides Bayesian posterior distributions of parameters for the model that may be used to construct point estimates of allele ages, as well as probabilities of population of origin, and of recurrent mutation. The model is designed for use with alleles that may be in high frequency and are not ascertained; it uses a sample of neutral “auxiliary” loci to extract information about relevant parameters of indirect interest such as migration rates between populations and population divergence times, integrating over the uncertainties of these parameters when making inferences about a target mutation within a particular gene/locus. The method assumes free recombination between loci and no recombination within loci, assumptions likely to be satisfied for smaller genes containing a target mutation and a sample of auxiliary loci widely distributed across a genome. Simulations are used to validate the correctness and statistical performance of the method and it is applied to analyze a set of seven SNPs in genes that affect human skin pigmentation in three populations: an African population from Kenya, a Han Chinese population, and a European population from Great Britain that are all part of the Thousand Genomes Project database (Byrska-Bishop *et al*. 2022).

## Methods

### Bayesian inference method

Consider a population tree with migration between populations described by the multispecies coalescent model with migration (MSC-M) (Flouri *et al*. 2023). Given multilocus sequence data for samples of individuals from each population, our goal is to estimate the posterior probability density of the age and population of origin of all observed mutations at a particular locus conditional on the population tree topology. Let ***X*** = {*X*_*i*_}be the sequence data (multiple sequence alignments) for *L* loci. Let **Θ** be the vector of parameters of the species tree, including the population divergence times, ***τ***, the effective population sizes, ***θ***, and migration rates, ***M***. Let **G** = {*G*_*i*_}be the gene trees, including topologies, coalescence times, and migration events for the *L* loci. Let ***ω*** be the DNA substitution model parameters. Following Flouri et al. (2023), the joint posterior probability density of the model parameters is

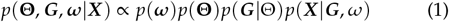

where *p*(***ω***) is the prior density of substitution model parameters, *p*(**Θ**) is the prior density of the MSC-M model parameters, *p*(***G*** | Θ) is the prior density of the gene tree given the MSC-M model, and *p*(***X*** | ***G, ω***) is the probability of the data (multiple sequence alignment) given the gene tree and substitution model parameters. We define the history of mutation *j* at locus *i* as a tuplet

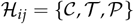

where 𝒞 ∈ {G, C, A, T} is the nucleotide created by the mutation, 𝒯 ∈ ℝ^+^ is the time (age) of the mutation, and 𝒫 is the population in which the mutation arose. Let ℋ_*i*_ be an array of mutation histories for locus *i*, then

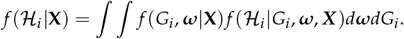

Although the mutation process is assumed to be Markovian, the individual mutations at a given site in a history do not occur independently of one another except under a simple Jukes Cantor DNA substitution model because the waiting time until a mutation occurs depends on the state, which is determined by the previous mutation. We will typically be interested in marginal of the mutation history variables, for example, *f* (𝒯_*ij*_ | ***X***) the posterior density of the time at which mutation *j* arose at locus *i* and

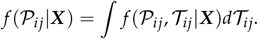

the posterior probability that mutation *j* arose in a particular population. We also will usually condition on 𝒞 (i.e., that the mutation produces a particular nucleotide). Two obvious difficulties are evaluating marginals for ℋ_*ij*_ and integrating over the posterior density of *G*_*i*_ and ***ω***. We solve this problem by using standard MCMC under the MSC-M model in the program bpp to generate samples from *f* (*G*_*i*_, ***ω*** | ***X***) and then simulate samples from *f* (ℋ_*i*_ | *G*_*i*_, ***ω, X***) conditional on each sample of the gene tree and substitution model parameters. During the MCMC, all of the parameters of the model are recorded for each sample, including gene trees with migration histories (***G***), substitution model parameters (***ω***), and MSC-M model parameters, (**Θ**). This simulation-based approach is similar to conventional Monte Carlo simulation except that the marginal samples from bpp are not strictly independent,

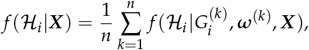

where the superscript *k* indicates the *k*th sample from the bpp MCMC analysis. By repeatedly simulating from 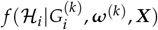 we approximate a sample from the density *f* (ℋ_*i*_ | ***X***). From these simulated mutation histories we can estimate the marginal posterior densities of the mutation history parameters. For example, if 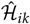 is the simulated history based on the *k*th MCMC sample the posterior probability that the mutation arose in population *a* is estimated as

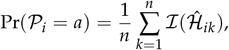

where 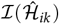 is equal to 1 if 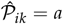 and 0 otherwise.

#### Simulating mutation historie

To simulate samples from 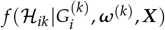, gene trees and the history of migration events on the gene trees are sampled from their posterior distribution using MCMC. Stochastic simulation (Nielsen 2002) is then used to simulate mutations on the sampled gene trees to generate a sample of the mutation times and populations of origin from the posterior distribution. For each gene tree sampled by the MCMC at a locus, we simulate mutational histories conditional on the data, the gene tree, and the substitution model parameters.

For a site *s* in the alignment, let *y*_*i*_ be the state at node *i* with daughter nodes *j* and *k*. Following Yang (2014) (eqn 4.5), let *L*_*i*_ (*y*_*i*_) be the conditional probability of the data of the tips descending from node *i* given the state at node *i* is *y*_*i*_.

For internal nodes in the tree, this is given by

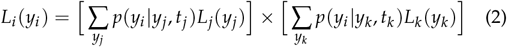

where *p*(*y*_*i*_ | *y*_*j*_, *t*_*j*_) is the transition probability of going from state *y*_*j*_ at node *j* to state *y*_*i*_ at node *i* in time *t*_*j*_ where *t*_*j*_ is the branch length connecting nodes *i* and *j*. For tips, *L*_*i*_ (*y*_*i*_) = 1 if *x*_*i*_ = *y*_*i*_ and 0 otherwise.

Let the root node have index *r* and *π*_*i*_ be the stationary frequency of base *i*. Then, we can calculate the probability for each root state as follows,

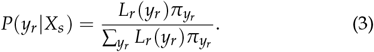

Given the state at the parent node and the data, we can also calculate the probability of each state at the daughter nodes. For a parent node *i* with daughter node *j*,

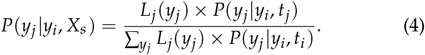

Eqns. 3 and 4 allow us to simulate the state of each node in a gene tree conditional on the sequence data and substitution model parameters using one of two possible methods. Mutation histories are then simulated along each branch on the gene tree conditional on the state at each node in the tree. If the states at the parent and daughter nodes are identical or the daughter node has missing data (or a gap), we use rejection simulation, simulating forward in time using the instantaneous rate matrix defined by the substitution model and its parameters. However, rejection simulation is inefficient when states of adjacent nodes require at least one mutation on a branch (i.e., the parent and daughter nodes differ). In that case, we instead simulate from the distribution of mutation times conditional on the end states of a branch and number of mutations. When end states differ, the minimum number of mutations is 1. Our focus is on closely related populations for which gene tree branches will typically be short (i.e., much less than 0.1 expected substitution per site). Thus, considering the probability of either 1 or 2 possible mutations on a branch closely approximates the total probability (the probability of 3 or more mutations is negligible). We derive the formulas for the generic case of a 4 state CTMC but then apply these to the specific case of HKY. Let *Q* be an instantaneous rate matrix for a 4 state CTMC satisfying 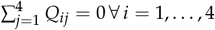. We denote *p*_*ij*_ (*t*) as the transition probability from state *i* to *j* on a branch of length *t*. We further define 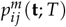 to be the transition probability from *i* to *j* on a branch of length *T* via *m* mutations occurring at times **t** = {*t*_1_, …, *t*_*m*_}. For *m* = 1 we have the probability

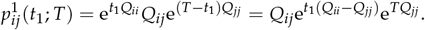

If we integrate out the time of the mutation we obtain the transition probability from *i* to *j* on a branch of length *T* via exactly 1 mutation,

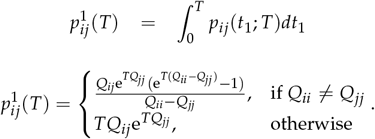

Now, given that a transition from *i* to *j* occurred on a branch of length *T*, the probability that it occurred via one mutation (versus one or more mutations) is

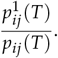

Conditional on one mutation the probability density of the time of the mutation is

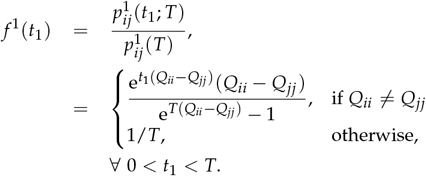

To simulate using the inverse transformation method (see below), we need the CDF

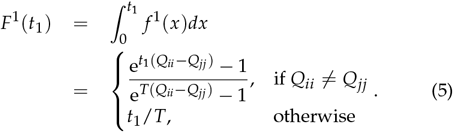

This is a conditional exponential distribution with rate *Q*_*ii*_ −\*Q*_*jj*_. We can also derive these probabilities for *m* = 2, although it is a bit tedious (Appendix).

This method of simulation assumes that *i* ≠ *j* and at most two mutations occur on any branch. Simulate *ξ* from a uniform distribution with PDF *f* (*ξ*) = 1, *ξ* ∈ (0, 1). If 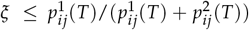 then simulate from *f*^1^(*t*_1_) otherwise simulate from 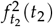. Since we are only interested in the time of observed mutations, we simulate from the marginal distribution of the time of the second mutation on branches with two mutations. We use the inverse transformation method to simulate from *f*^1^(*t*_1_). We find the inverse of Eqn. 5.

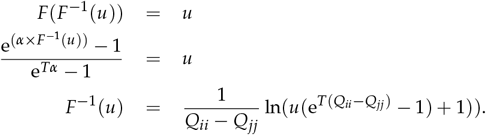

To simulate from 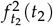 (Eqn. 6), we simulate *z* from a uniform distribution with PDF *f* (*z*) = 1, *z* ∈ (0, 1). Then we solve the equation 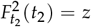 for *t*_2_ numerically using the bisection method.

#### HKY model

Let *p*_*ij*_ (*t*) be the transition probability for a HKY model with transition and transversion rates *α* and *β* and stationary frequencies *π*_*T*_, *π*_*G*_, *π*_*C*_ and *π*_*A*_ = 1 − *π*_*T*_ − *π*_*G*_ − *π*_*C*_. The probability of a mutation from nucleotide A to T on interval (0, *T*) via exactly one mutation at time *t* is

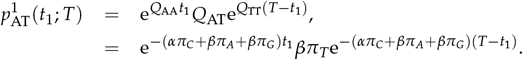

Other probabilities can be obtained by substitution in a similar way. For the HKY model, there are analytical results for transition probabilities *p*_*ij*_ (*t*) (Yang 2014). We can find the probability that the transition from A to T on a branch of length *T* occurred via one mutation,

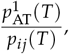

or two mutations,

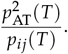

The sum of these probabilities typically is near one for *T* ≪ 0.1, assuming branch length is in units of expected substitutions. We can check the absolute error of the two mutation approximation by calculating the residual probability,

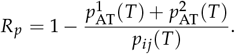

The residual should ideally be very small. If such is the case, allowing either one or two mutations per branch in our simulations should provide sufficient accuracy. Under a Jukes Cantor model, for example, with T = 0.1, *R*_*p*_ = 0.001283.

#### Summary of inference results

For each mutation that has an observed descendant in the tree, the time of the mutation and the population in which the mutation occurred is recorded. Repeating the simulation procedure for all samples of the MCMC and for each variable site generates a posterior distribution of the mutation time and population of origin for each variable site.

### Simulation Study

#### Validity and accuracy of posterior probabilities

To assess the correctness of the program, datasets were simulated with five sets of simulation conditions (Table S1). All migration rates not listed in Table S1 were set to zero. For each set of conditions, 50 replicate datasets were simulated using the simulation method implemented in bpp. 5 samples of 100 loci were collected from each population. Loci were 5000 bp in length. For simulations 1 and 2, an HKY model was used with *κ* = 2 and base frequencies *π*_*A*_ = 0.24, *π*_*C*_ = 0.26, *π*_*G*_ = 0.23, *π*_*T*_ = 0.27. For simulations 3 - 5, an HKY model was used with *κ* = 4 and base frequencies *π*_*A*_ = 0.2, *π*_*C*_ = 0.3, *π*_*G*_ = 0.23, *π*_*T*_ = 0.27. A strict clock was used for all simulations.

For each dataset, inference was performed with bpp using an HKY model with a strict clock. The root age prior was assumed to be a gamma distribution with *α* = 5 and *β* = 50, 000, denoted as Γ(5, 50000). The *θ* prior was Γ(2, 2000). The migration rate prior was Γ(10, 500) for simulations 1 and 3, Γ(1.5, 15) for simulations 2 and 4, and Γ(1.5, 3) for simulation 5. The same migration connections were included in the inference as were used in simulation for each analysis. For each dataset, two independent MCMCs were run. The MCMCs were sampled every 8 iterations, for a total of 250,000 samples (2 × 10^6^ iterations) with a burnin of 100,000 for simulations 1-4. For simulation 5, the total number of samples was doubled as mixing was worse than in simulations 1-4.

MCMC convergence was checked by comparing the results from independent pairs of MCMCs using a two-sample t-test,

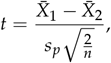

where

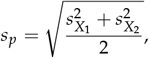

*n* is the MCMC iteration sample size, *X*_*i*_ are the sample means and 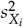 are unbiased estimates of the variance (Nagel *et al*. 2024). Samples from MCMCs are not independent, so we let *n* = 2, 000 rather than the actual number of samples in the MCMC. This test was performed on all *τ*s, *θ*s and *M*s in the model. If any of the parameters were significantly different between the two MCMCs or the effective sample size was less than 200 for any parameter, the pair of MCMCs was manually compared using tracer (Rambaut *et al*. 2018). Of the pairs manually inspected, almost all had very similar distributions for all parameters, suggesting the convergence criteria were conservative. Only one pair of MCMCs had substantially different distributions. This pair was part of simulation 5, and these replicates were removed from subsequent analyses. All other manually inspected MCMCs appeared to have converged.

#### Statistical performance of method

The statistical performance of the method under different conditions was studied by simulating data with varying sequence lengths or numbers of samples. For all simulations, bpp was used to simulate gene trees and sequence data using the 3 population tree shown in Figure 1a. Parameters values for mutation scaled contemporary and ancestral effective population sizes were *θ*_*A*_ = 4.9 × 10^−4^, *θ*_*B*_ = 4.4 × 10^−4^, *θ*_*C*_ = 2.1 × 10^−3^, *θ*_*AB*_ = 8.7^−5^, *θ*_*ABC*_ = 6.1 × 10^−4^, and for divergence times were *τ*_*AB*_ = 2.4 × 10^−5^, *τ*_*ABC*_ = 4.7 × 10^−4^. The migration rate parameter values were *M*_*A* → *B*_ = 0.15, *M*_*B* → *A*_ = 0.05, *M*_*A* → *C*_ = 0.12, *M*_*C* → *A*_ = 0.1, *M*_*B* → *C*_ = 1.5, *M*_*C* → *B*_ = 0.36, *M*_*AB* → *C*_ = 5.0, *M*_*C* → *AB*_ = 0.61. Sequence data were simulated on gene trees using a strict clock and an HKY model with parameters *π*_*A*_ = 0.28, *π*_*C*_ = 0.23, *π*_*G*_ = 0.22, *π*_*T*_ = 0.27, and *κ* = 8.0. These parameters were chosen to be similar to the estimates for the empirical dataset.

**Figure 1.**
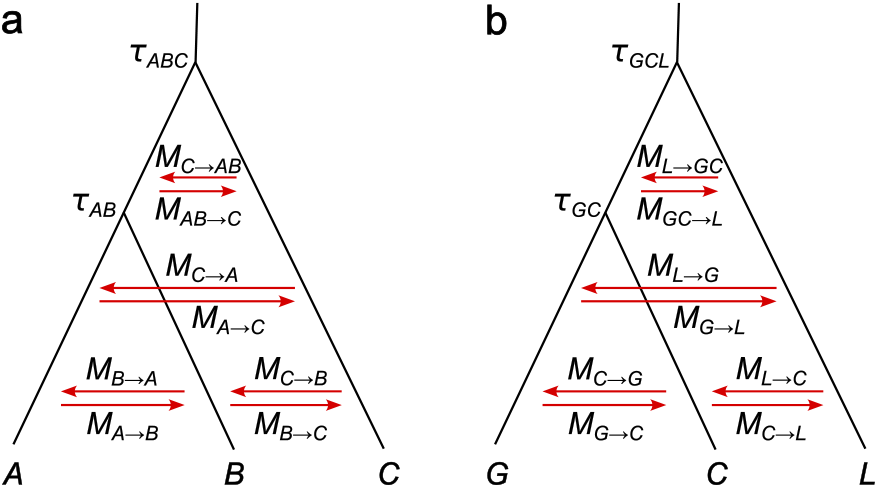
Trees for the simulation and empirical analysis. (a) The population tree used for the simulation study on the impact of locus length and number of samples. (b) The population tree for the analysis of the human skin pigmentation dataset. The first letter of the population named is used in the figure. *G* is for GBR, *C* is for CHS and *L* is for LWK.

In the simulations, mutation age and population of origin inferences were only conducted on 10 loci. These are referred to as the focal loci. Auxiliary loci were simultaneously analyzed in bpp to precisely estimate MSC parameters (*τ*s, *θ*s, and *M*s). For all simulated datasets, 500 auxiliary loci of length 5000 bp were simulated, each with 20 sequences per population. We conducted 30 replicates simulations for each set of simulation conditions.

It is possible to have multiple mutations segregating at a simulated site. However, the focus of this study is on sites that have a single mutation since the properties of the method are more difficult to summarize when there are multiple mutations at a site and this outcome is relatively rare under the conditions we consider. Sites with multiple mutations were therefore excluded from the results. The focal loci may have multiple mutations, but the mutation age and population of origin results only include the first mutation in the alignment that meets the inclusion criteria.

Given a mutation of interest, different lengths of adjacent sequence could be used in the analysis. Longer sequences should contain more information about gene tree topology and branch lengths. However, exceptionally long sequences may violate the assumption of no recombination within a locus. Focal sequences of length 10 kb were simulated. Two smaller alignments of length 1 kb and 5 kb were created from each 10 kb alignment. The smaller alignments began at the first mutation; additional sites were added until the desired sequence length was reached. If the end of the alignment is reached before achieving the desired length, sequence is appended from the beginning of the alignment. Since recombination was not simulated and sites in the alignment are independent, it does not matter that the sequence used in the analysis was not continuous in the original alignment.

The number of samples (diploid individuals) at a locus may impact the inferences. The focus of our simulations was on common mutations; such mutations are likely to be observed in a random non-ascertained sample. For each population, 1000 sequences of length 5000 bp were simulated at each of 10 focal loci. We conditioned on a mutation being present in at least 25% of the samples in at least one population. At least one mutation met this condition in all simulations, and no simulations were rejected. From the sample of 1000 sequences, either 10, 100, or 200 sequences were subsampled at each locus in each population. If a subsample did not contain both variants, the sample was redrawn. This occurred only three times, all with samples of size 10. The subsampled sequences were chosen independently which allows the same sequence to appear in different subsamples.

Each simulated dataset was analyzed using bpp with the true population tree and a saturated migration model. An HKY substitution model with a strict clock was used. The root age prior was Γ(5, 50000), the *θ* prior was Γ(2, 2000), and the *M* prior was Γ(1.5, 3). For each dataset, two MCMCs were run with a burnin length of 100,000, sampling every 8 iterations for a total of 500,000 samples. Convergence was checked as described above, but with *n* = 200. The simulations that were checked by hand had very similar distributions and no datasets were removed. MutAnce was run for the focal loci in all analyses on one MCMC output from each pair. To investigate the best-case scenario with known gene trees (infinite data), MutAnce was run with the true gene tree, migration history, population split times, and substitution model parameters. The true migration history is in reality never known, even with sequences having an infinite number of sites, but this procedure still provides a conservative upper bound on method performance.

### Analysis of human skin pigmentation SNPs

To infer the time and population of origin of mutations impacting skin pigmentation in humans (Table 1), a dataset composed of both (focal) loci (flanking SNPs influencing skin pigmentation) and random (auxiliary) loci, was constructed using data from the 1000 Genomes Project (TGP). The random loci provide information that refines estimates of MSC parameters from bpp, including migration rates, population divergence times, and effective population sizes. One population from each of three continents was used: Luhya from Kenya (LWK), British from England and Scotland (GBR), and Southern Han Chinese (CHS) from South China. Phased VCF files from the TGP and a GRCh38.p14 reference genome and annotations (release GCF_000001405.40-RS_2023_10) were downloaded. Children of TGP trios were excluded.

**Table 1.** Skin pigmentation SNPs.

| Gene | SNP ID | SNP coordinate | gene coordinates | sequence coordinates in analysis |
| --- | --- | --- | --- | --- |
| TYR | rs1042602 | 11:89178528 | 11:89177875-89295759 | 11:89177875-89182874 |
| ASIP | rs6058017 | 20:34269192 | 20:34186493-34269344 | 20:34264345-34269344 |
| OCA2 | rs1800404 | 15:27990627 | 15:27719008-28099315 | 15:27988127-27993126 |
| MATP | rs16891982 | 5:33951588 | 5:33944623-33984693 | 5:33949088-33954087 |
| MC1R | rs2228479 | 16:89919532 | 16:89918862-89920972 | 16:89918862-89920972 |
| MC1R | rs2228478 | 16:89920200 | 16:89918862-89920972 | 16:89918862-89920972 |
| SLC24A5 | rs1426654 | 15:48134287 | 15:48120990-48142672 | 15:48131787-48136786 |

To choose auxiliary loci, coordinates for all exons in each of chromosomes 1-10 were obtained from reference sequence annotations. Many exons overlap and coordinates were combined within overlapping exons. All exon regions less than 5 kb in length or within 20 kb of the closest exon sequence in the list of pigmentation SNP loci were removed. Of the remaining exon coordinates, 110 were randomly chosen from each of chromosomes 1-10 and coordinates were obtained for the initial 5 kb of each for use as auxiliary loci in the analysis (independent of whether an exon is transcribed from the plus or minus strand). For auxiliary loci 10 individuals were randomly chosen from each population. The sequences for both phased haplotypes were extracted from the VCF file for each individual using bcftools with consensus and the options –regions -overlap 2 -H 1p 1U -s and –regions -overlap 2 -H 2p 1U -s for the first and second haplotypes (Li 2011). The sequences were aligned using mafft (v7.453) with the default parameters (Katoh *et al*. 2002). Auxiliary loci with more than 200 gaps for any individuals were removed from the set of 110 loci for that chromosome. Of the remaining loci, 100 were used for further analysis.

For focal loci, sequence coordinates were determined by the location of the candidate pigmentation SNP in the gene and the length of the gene. If the gene was less than 5 kb, the entire gene was used in the analysis. If a SNP was in the first (or last) 5 kb of the gene, the first (or last) 5 kb was used in the analysis, respectively, otherwise the region starting 2500 bp before and terminating 2499 bp after the SNP was used. For the focal loci (Table 1) 20 individuals were randomly chosen from each population. Both phased haplotypes for the 60 individuals were extracted from the VCF files using bcftools, resulting in 40 sequences per population for each skin pigmentation locus. The sequences were aligned using mafft. There were no large deletions in the alignments, and the loci were not within 20 kb of any of the randomly selected loci.

The skin pigmentation loci were combined with the randomly selected loci to create a dataset of 1006 loci that was analyzed using bpp with CHS and GBR as sister populations and LWK as the outgroup (Fig. 1b). A HKY model with a strict clock was used. The prior on root age was Γ(2.5, 40000) and the prior on *θ* was Γ(2.5, 5000). A saturated migration model was used with migration connections between all possible pairs of contemporary and ancestral populations (Fig. 1b). and migration rate prior Γ(1.5, 3). The burnin for the MCMC was 200,000 iterations and the MCMC was sampled every 8 iterations for a total of 500,000 samples. Three MCMCs were run and convergence was assessed by comparing the posterior distributions from the three runs in tracer (Rambaut *et al*. 2018). For the skin pigmentation loci, MutAnce was run only on the SNP of interest. To convert between units of expected number of mutations and years, a mutation rate of 10^−9^ mutations per year was assumed (Scally and Durbin 2012).

## Results

### Simulation study

#### Validation and accuracy of posterior probabilities

MutAnce was run on every locus for each of the 50 replicates in simulations 1-5. The inferences were summarized across all loci and replicates within a simulation set. The coverages (probability the true time was contained in the 95% credible interval) for the times were 0.947, 0.947, 0.948, 0.948, 0.950 for simulations 1-5, respectively. To assess whether the posterior probabilities of the population of origin and the derived base were accurate, the posterior probabilities for each were placed into bins ([0-0.1), [0.1-0.2), …, [0.9-1.0), [1.0]). For each bin, the average posterior probability and the proportion correct were calculated. For all of the simulations 1-5, the average posterior probability in each bin closely matched the proportion of correct inferences in the bin (Fig. 2). The summary results only include cases with a single mutation at a site since this is the focus of our study, and there were relatively few cases of multiple mutations in the simulated datasets.

**Figure 2.**
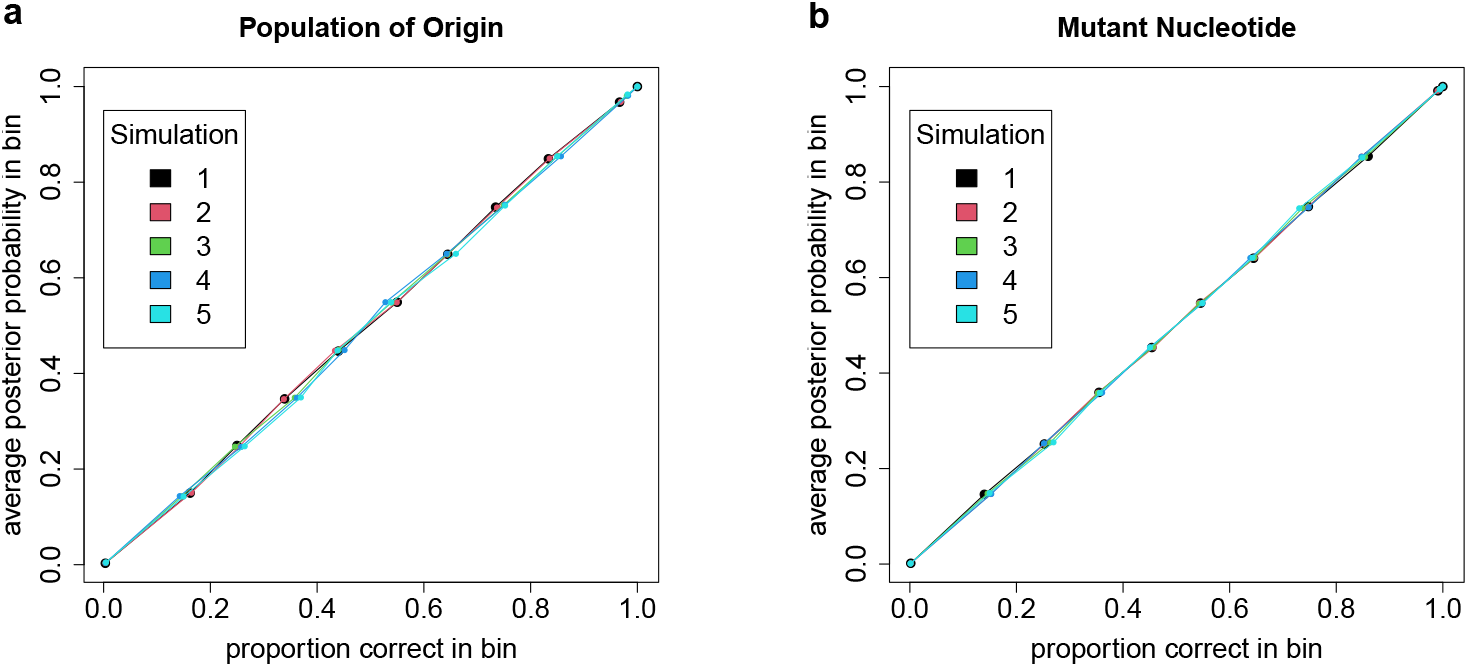
Results obtained from analyses of simulated sequence data using the BPP and Mutance programs to infer posterior probabilities of population of origin of a mutation and derived nucleotide state. The average posterior probability in each bin is linearly related to the proportion correct in each bin for the population of origin (a) and mutant nucleotide (b).

#### Statistical performance

The coverage for the time estimates is close to 95% independent of the locus length (Fig. 3a). The bias is negative (underestimating mutation age) and decreases as the locus length increases, but the absolute value increases (Fig. 3b). The bias with a sequence length of 10,000 bp is similar to that obtained by assuming the gene tree is known (infinite sequence length). The root mean square error (RMSE) and the size of the 95% credible intervals decrease with decreasing sequence length (Fig. 3b, S1 a). With infinite sequence length (known gene trees), the uncertainty in the mutation time remains relatively large, on the order of 25,000 years assuming a mutation rate of 10^−9^ per year.

**Figure 3.**
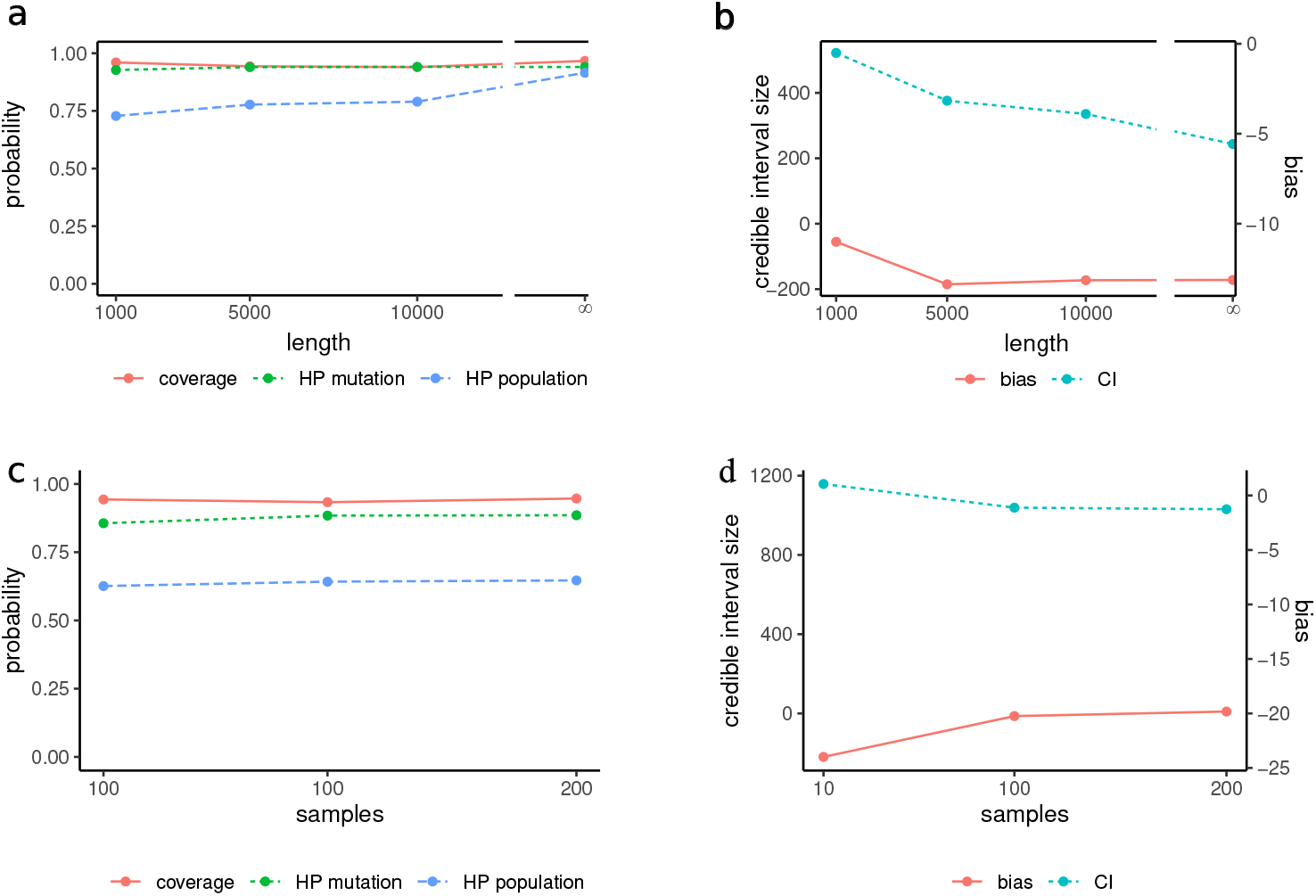
Summary statistics based on locus length and number of samples. Panels (a) and (c) show the probability the true time in the 95% credible interval (coverage) (red), the probability of the mutation with the highest posterior density (green), and the probability of the population with the highest posterior probability (blue), for the locus length and sample size simulations, respectively. Panels (b) and (d) show the size of the credible interval in ka (teal) and bias in ka years (red) for the locus length and sample size simulations, respectively. The infinite point corresponds to knowing the true gene tree with migration events and the true species divergence times. For all plots in ka, a mutation rate of 10^−9^ mutations per year was assumed.

There is considerable uncertainty about the population of origin (Fig. 3a, 4a), but less uncertainty about the mutant nucleotide (Fig. 3a, 4b). Increasing the sequence lengths leads to higher posterior probability for the correct population in some, but not all, cases. Even assuming a known gene tree, including migration events, and known population divergence times, population of origin inferences remain uncertain. Inferences for mutations with a true origin in population AB appear to improve most in this case.

**Figure 4.**
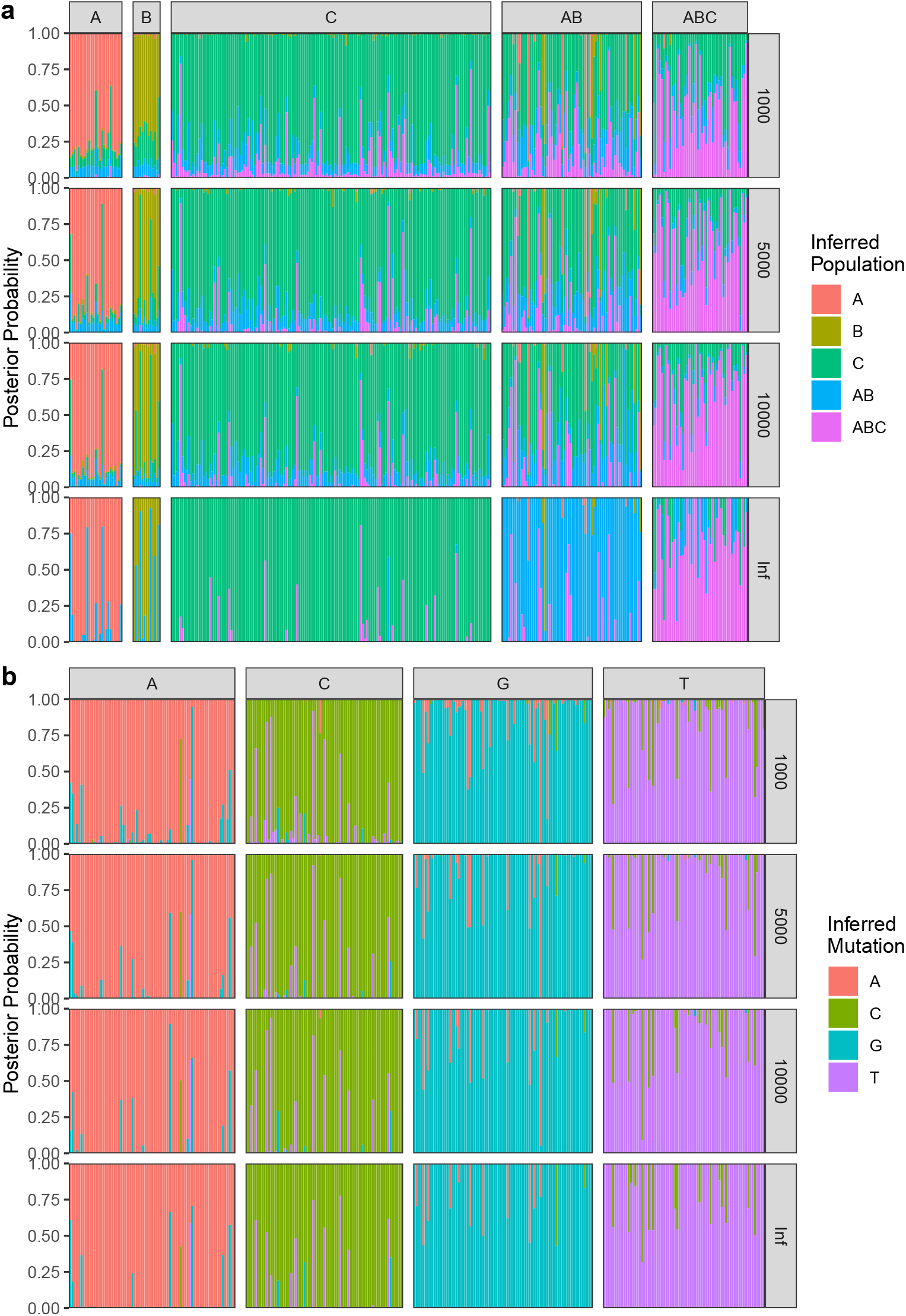
The posterior probability associated with each (a) population of origin and (b) mutation for the locus length simulations. On the x-axis, each bar shows the posterior probability for a single mutation. The true populations and mutations are shown across the top while the inferences are shown by the color. The locus length increases from top to bottom.

In general, the average performance of the method is minimally impacted by the number of samples (Fig. 3c,d). As the number of samples increases, the mean bias increases and becomes positive (Fig. 3d) and the RMSE and size of the 95% credible intervals decrease slightly (Fig. 3d, S1 b). The coverage and the posterior density of the highest posterior populations and mutations do not change appreciably based on the number of samples (Fig. 3c).

In the sample size simulations, there were no mutations that occurred in populations A or B that met the inclusion criteria of being present in at least 25% of the samples in at least one population. The mutations were older on average for sample size simulations (760 ka) than the locus length simulations (450 ka). The population of origin and the mutations were more uncertain in the sample size simulations (comparing Fig. 3d to Fig. 3c and Fig. 4 to Fig. 5).

**Figure 5.**
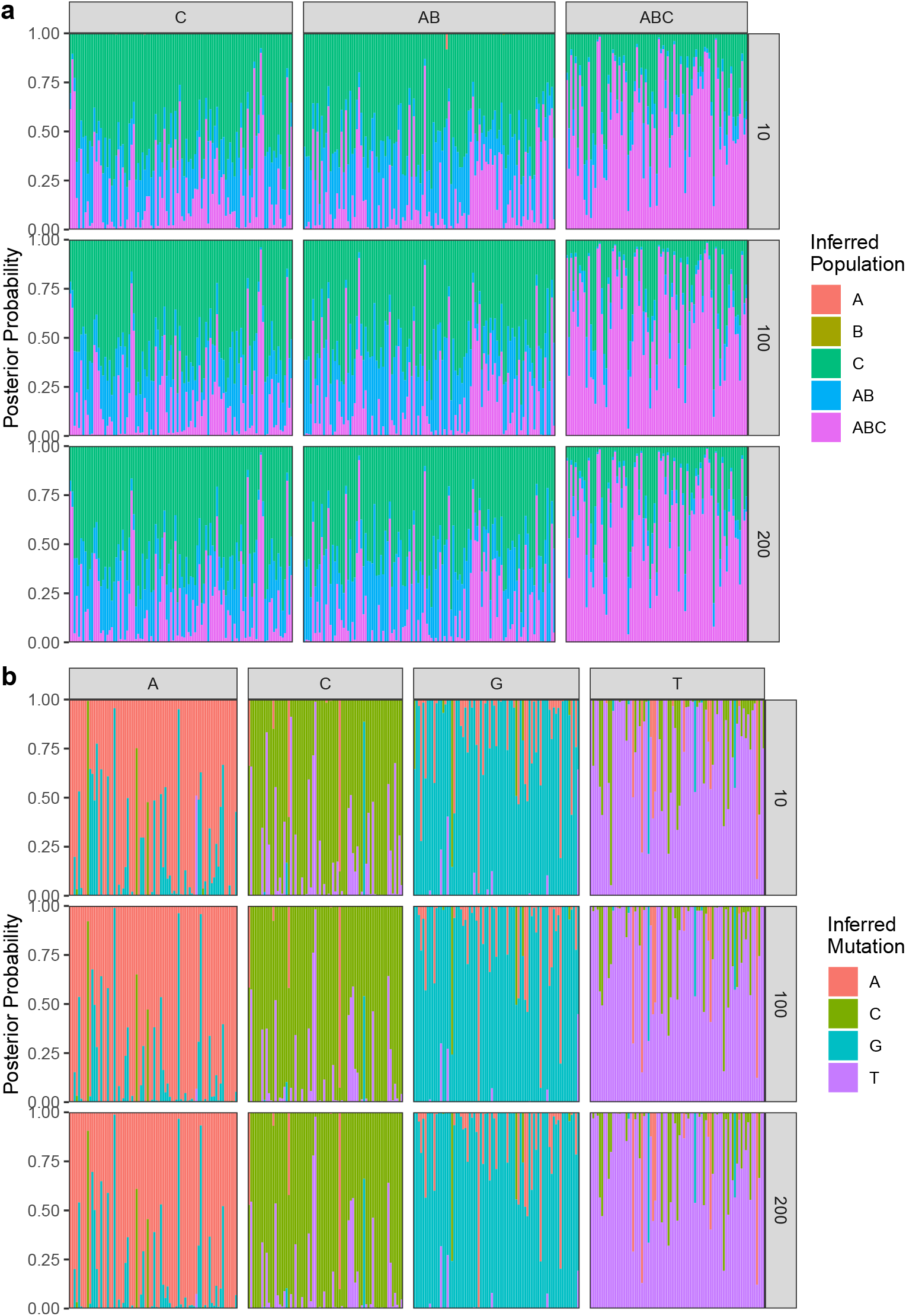
The posterior probability associated with each (a) population of origin and (b) mutation for the sample size simulations. Each bar on the x-axis shows the posterior probabilities associated with a single mutation. The true populations and mutations are shown across the top while the inferences are shown by the color. The number of samples increases from top to bottom.

### Empirical results: skin pigmentation

The putative ancestral allele for the skin pigmentation alleles has high posterior probability (> 0.99) for all SNPs except for one of the focal SNPs in MC1R (rs2228478) (Table 2). For the focal SNP in OCA2, there were multiple mutations with high probability (> .99). For the focal SNPs in MC1R (rs2228478) and MATP, the posterior probabilities of multiple mutations were intermediate. All the other SNPs had a single origin with high posterior probability (> 0.99). The posterior probability of recurrence (a mutation to the same base occurs multiple times) is non-negligible for all of the SNPs with a non-negligible probability of multiple mutations. For the focal SNP in MATP, the probability of recurrent mutations given there are multiple mutations is close to one, whereas mutations to the ancestral state are more probable for the focal sites in OCA2 and MC1R (rs2228478).

**Table 2.** Inference of mutation history for skin pigmentation alleles.

| SNP | putative ancestral allele | ancestral allele (marginal probability) | P(multiple mutations) | P(recurrent) |
| --- | --- | --- | --- | --- |
| rs1042602 | C (Olalde *et al.* 2014) | C (1.00) | 0.0079 | 0.00 |
| rs6058017 | G (Bonilla *et al.* 2005) | G(1.00) | 0.0068 | 0.00 |
| rs1800404 | C (Crawford *et al.* 2017) | C (0.99) | 0.9997 | 0.73 |
| rs16891982 | C (Olalde *et al.* 2014) | C (1.00) | 0.1724 | 0.17 |
| rs2228479 | G (Olalde *et al.* 2014) | G (1.00) | 0.0061 | 0.00 |
| rs2228478 | G (Olalde *et al.* 2014) | G (0.5247) | 0.8033 | 0.46 |
| rs1426654 | G (Crawford *et al.* 2017) | G (0.995) | 0.0085 | 0.00 |

The probability of recurrent is the probability that a mutation to the same state (base) occurs twice conditional on two or fewer mutations. The conditioning only impacts the results of rs2228478, where the probability of three or more mutations is 0.0109, which is larger than all other SNPs.

The uncertainty in the mutation ages tends to be large, especially for the mutations that are inferred to be older (Fig. 6). The uncertainty in the age of the mutations is much larger than the uncertainty in the population divergence times. Thus the uncertainty in the population of origin is not driven by the uncertainty in the population age, but is probably due to the uncertainty in the migration histories and the mutation ages.

**Figure 6.**
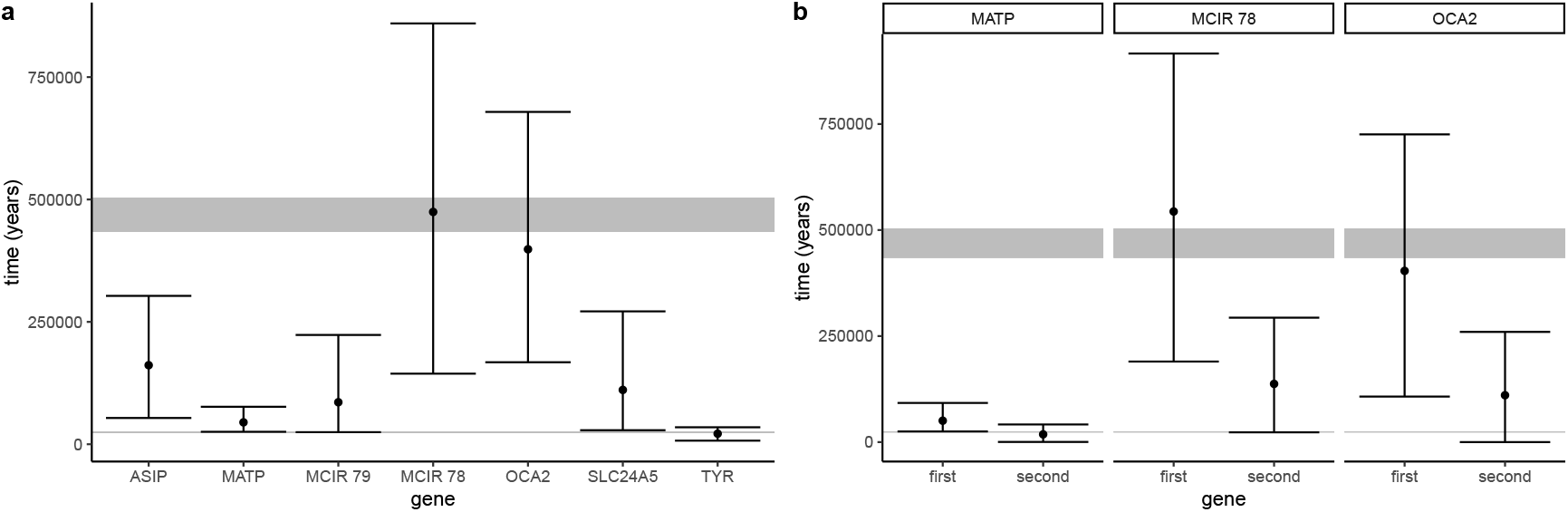
The inferred time of the mutation in each skin pigmentation gene conditional on their being one (a) or two (b) mutations at the site. The points show the mean and the bar shows the 95% highest posterior density interval. The gray bars show the 95% credible intervals population divergence times. The thin gray bar close to 0 is the divergence between CHS and GBR. The larger gray bar is the divergence between LWK and the common ancestor of GBR and CHS. Time zero is time present, so larger values are older. In (b), first and second refer to the first and second mutations that occurred in forward time. MC1R 79 and MC1R 78 refer to rs2228479 and rs2228478, respectively. A mutation rate is assumed to be 10^−9^ mutations per base per year.

The mutations in ASIP, MATP, and MCIR (rs2228479) are inferred to have occurred in either the common ancestor of GBR and CHS, or in LWK, with high posterior probability (Fig. 7). The focal mutations in MC1R (rs2228478) and OCA2 are inferred to occur in those populations or the common ancestor of all populations. None of the mutations are inferred to have occurred in CHS and only TYR has a substantial posterior probability of having occurred in GBR conditional on a single mutation. However, conditional on two mutations, there is substantial posterior probability associated with the second mutation at the focal site in MATP occurring in GBR.

**Figure 7.**
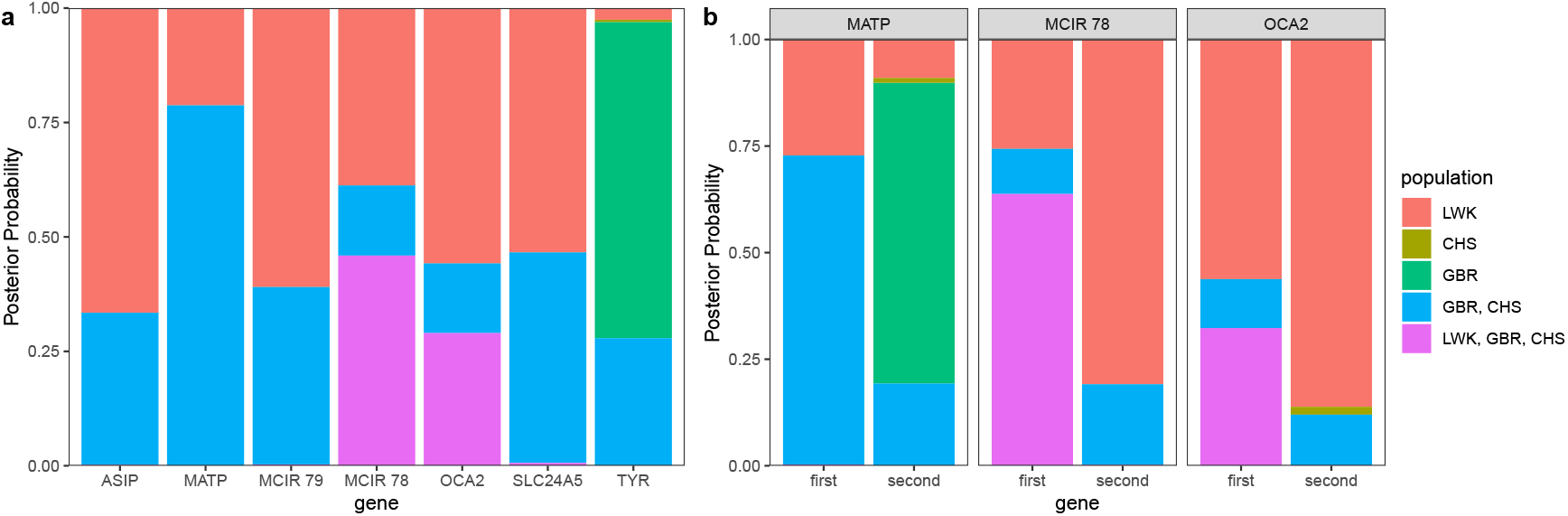
The inferred population of origin of the mutation in each skin pigmentation gene conditional on there being one (a) or two (b) mutations at the site. MC1R 79 and MC1R 78 refer to rs2228479 and rs2228478, respectively.

## Discussion

A new method was developed to estimate the time and population of origin of a mutation in structured populations and was implemented in the program MutAnce. The method also infers the ancestral and derived states for the site. The method performs well, with accurate posterior probabilities and credible intervals for the times, populations of origin, and the ancestral and derived states. The estimates of time and population of origin often have considerable uncertainty for simulated and empirical datasets, while ancestral and derived states are more precisely estimated. This uncertainty persists even when all parameters of the model are known, suggesting the uncertainty largely results from the stochasticity in the mutation process. For example, in the simple case of a JC model, given different base states at the ends of a branch in the gene tree, the probability distribution for the time of the mutation is uniform. Conceptually, adding samples may subdivide branches in the gene tree, potentially narrowing the possible time interval on a branch during which a mutation could have occurred. However, in our simulations, increasing the number of samples has minimal impact on the average size of the 95% credible intervals, RMSE, and coverage for the estimates of time. This is likely because additional samples tend to coalesce quickly, so the number of lineages persisting deeper in the gene tree is not greatly altered. This leads to long branches deeper in the gene tree and uncertainty in the estimates. Increasing locus length improved inferences, but the improvement was greater from 1000 to 5000 bp than 5000 to 10,000 bp. This suggests that intermediate length loci might be ideal to increase the power of the method while reducing the chance of within locus recombination. While our simulations did not include recombination, the assumption of no recombination within a locus is expected to be violated more often as the locus length increases.

The method was used to analyze seven SNPs associated with human skin pigmentation in three populations (Asian, African and European). The SNPs occur in six protein coding genes: ty-rosinase (TYR), agouti signaling protein (ASIP), oculocutaneous albinism II (OCA2), membrane associated transporter protein (MATP), melanocortin-1 receptor (MC1R), and solute carrier family 24 member 5 (SLC24A5). It is unclear how to infer the age of these mutations using classical methods to make comparisons since they occur in multiple populations. For simplicity, we treated each population as independent when applying these older methods, which sometimes resulted in very different estimates of age for the same SNP in different populations (Table S1).

The protein encoded by the TYR gene catalyzes the first two steps in the melanogenesis pathway (Sturm *et al*. 2001). This gene is associated with skin color and tanning ability and when mutated is the cause of almost half of albinism cases in Caucasians (Rooryck *et al*. 2009). The SNP rs1042602 in TYR is associated with skin color and susceptibility to melanoma and squamous cell carcinoma (Nan *et al*. 2009). The putative derived allele, A, is associated with light skin and eye color and the lack of freckles. The derived allele A is at high frequency in Europeans, and the putative ancestral allele, C, is at high frequency in Africans. Other SNPs in TYR are also associated with cutaneous melanoma and basal cell carcinoma (Gudbjartsson *et al*. 2008). Two different methods have been used previously to estimate the age of the derived TYR allele A. One estimate was 6.1 ka and the other was 15.6 ka (Hudjashov *et al*. 2013). It is speculated that TYR has been under strong selection in Europe starting 11 to 19 ka (Beleza *et al*. 2013), and continuing until at least 5 ka (Wilde *et al*. 2014). Our inferred age of the allele (22 ka, 95% credible interval 7.3-35 ka) is older than previous age estimates, which is consistent with standing genetic variation existing prior to selection. The derived allele has a high posterior probability of having arisen in either the European (GBR) or ancestral Eurasian population (GBR,CHS) and very low probability of having arisen in the African (LWK) population

ASIP is a major determinant of pigmentation in mice (Suzuki *et al*. 1997). SNPs in this gene are also associated with cutaneous melanoma and basal cell carcinoma (Gudbjartsson *et al*. 2008; Nan *et al*. 2009). The SNP rs6058017 in ASIP is in the 3’ untranslated region. The putative ancestral G allele is associated with dark hair and brown eyes while the putative derived A allele is associated with lighter hair and eye colors in European Americans (Kanetsky *et al*. 2002). The G allele is also associated with darker skin pigmentation in Africans (Bonilla *et al*. 2005). Our inferred age of the A allele is 160 ka (95% credible interval 54 - 300 ka) which places the time of origin of this allele between the divergence of all three populations and the divergence of GBR and CHS. There is a high posterior probability that the derived A allele arose in either the African population (LWK) or the ancestral Eurasian population (GBR,CHS).

OCA2 (formerly P gene) is a chloride transporter which impacts pigmentation by changing the pH in melanosomes (Bellono *et al*. 2014). Mutations in this gene are the most common cause of albinism in Africans (Brilliant 2001), and the gene is associated with skin, hair, and eye color (Cook *et al*. 2009). The putative ancestral variant of rs1800404, C, is associated with dark pigmentation and is common in Africans, South and East Asians, and Austrlo-Melanesians. The putative derived allele, T, is common in Europeans (Crawford *et al*. 2017). Some have suggested that this locus experienced balancing selection. The time of the most recent common ancestor (tMRCA) of the T allele was estimated by Crawford *et al*. (2017) to be 629 ka (95% confidence interval 426-848 ka). We inferred there to be multiple mutations at this site with high posterior probability (>.99) and likely recurrent mutation (posterior probability >.70). The first mutation is slightly younger than the previous estimate of the tMRCA of the T allele (400 ka, 95% credible interval 110 - 730 ka), while the second mutation appears younger (220 ka, 95% credible interval 0.001 - 260 ka). The population of origin of the derived allele(s) has a high posterior probability of being either African (LWK), the Eurasian population ancestor (GBR,CHS) or an a populations ancestral to all three populations (LWR,GBR,CHS).

MATP, also known as solute carrier family 24 membrane 2 (SLC45A2), is a membrane associated transported protein that plays a role in the distribution and processing of tyrosinase as well as other pigment related enzymes (Costin *et al*. 2003). This gene is associated with hair, skin, and eye color and tanning ability (Graf *et al*. 2005; Nan *et al*. 2009). The putative derived allele, G, of SNP rs16891982 is associated with light skin, hair, and eye color while the alternative allele, C, is associated with darker color (Norton *et al*. 2006; Cook *et al*. 2009). Strong selection on this SNP is estimated to have started 11-19 ka ago in Europeans (Beleza *et al*. 2013). This is one of the younger alleles in our dataset and the age is inferred to be 45 ka (95% CI 25 - 76 ka, conditional on a single mutation). It is possible there were multiple mutations at this site (posterior probability .17), but the time of origin is older than the previously inferred time of selection independent of the number of mutations. If a second mutation recurred it very likely to have originated in the European population (GBR).

MC1R, melanocortin-1 receptor, regulates the ratio of pheomelanin and eumelanin (Valverde *et al*. 1995). It is involved in animal coat color in many mammals and there has been a rapid accumulation of polymorphism in many vertebrates (Switonski *et al*. 2013; Hofreiter and Schöneberg 2010). This gene is associated with pheomelanic red hair, fair skin, freckling, tanning, and skin cancer risk (Beaumont *et al*. 2009). Many variants are classified as either high or low penetrance. High penetrance variants are associated with with basal cell carcinoma (Kosiniak-Kamysz *et al*. 2012). We analyzed two SNPs at this locus, rs2228479, also known as V92M, and rs2228478. The SNP rs2228479 is low penetrance and the A allele is associated with red hair and fair skin (Valverde *et al*. 1995). The Altai Neanderthal carried the V92M allele, and some authors have suggested this allele is of Neanderthal origin (Ding *et al*. 2014). Our estimate of the age (85 ka, 95% credible interval 25 - 220 ka) allows the mutation to have occurred in humans or Neanderthals. The SNP rs2228478 is synonymous substitution and is in linkage disequilibrium with V92M. Though included as a candidate SNP in association studies, it is not generally found to be associated with pigmentation phenotypes (Mengel-From *et al*. 2009; Stokowski *et al*. 2007; Cao *et al*. 2013).

The gene SLC24A5 gene determines the golden pigmentation phenotype in zebrafish (Lamason *et al*. 2005). In mice, mutations in this gene are associated with smaller and paler melanosomes and less overall pigmentation (Vogel *et al*. 2008). In humans, it is associated with skin color, as is the SNP rs1426654 (Quillen *et al*. 2012; Norton *et al*. 2006; Crawford *et al*. 2017). The A allele is common in European, Pakistani, and Indian populations while the G allele is common in West Africa and East Asian populations (Crawford *et al*. 2017). The estimated tMRCA of the lineages containing the A allele has been estimated to be 29 ka. It has been estimated that this allele was introduced into East African more than 5 ka and has since risen to high frequency, possibly as a result of selection (Crawford *et al*. 2017). The estimated tMRCA of the lineages containing the putative derived A allele is much younger than our allele age estimate (110 ka, 95% credible interval 29 - 270 ka). This may be due to the inclusion of the CHS population in our dataset and a structured population analysis. There is evidence of a near complete selective sweep in European populations; the gene is in a large continuous region of low heterozygosity in European genomes (Lamason *et al*. 2005). This sweep may have acted on standing genetic variation as our analysis suggests that the derived allele very likely arose in either the African population (LWK) or the Eurasian ancestral population (GBR,CHS).

The results of our analyses agree with many previous inferences of ancestral allele states and our simulations suggest that ancestral state can be inferred confidently, suggesting that many existing inferences on ancestral state are accurate. In this dataset, the mutations tended to be old, with most of them occurring before the divergence between GBR and CHS and two of the mutations possibly occurred before the population divergence between GBR/CHS and LWK, which corresponds to an ancestral population in Africa before the out of Africa event. Overall, there is considerable uncertainty concerning the population in which those mutations occurred, with our results often supporting multiple possibilities including LWK, the common ancestor of GBR and CHS, or the common ancestor of all three populations. This uncertainty about population of origin is likely driven by uncertainty in the mutation age – not the divergence times between the populations – because mutation age is much more uncertain than divergence times. By contrast, there is strong evidence that none of the putative derived mutations that we considered arose in the Asian (CHS) population.

There is strong evidence of multiple mutations in some cases (the focal mutation OCA2 and the SNP rs2228478 in MC1R). This could be an artifact of recombination within the locus. There are no programs that jointly infer recombination and mutation; either recombination is inferred assuming no mutation or mutation is inferred assuming no recombination (Rannala and Slatkin 1998). However, this could be investigated by varying the sequence length and the break points in the sequence. Varying the size and position of the window of sequence around the mutation may remove recombination events effecting the locus. Alternatively, future work could develop a method which jointly infers mutation and recombination. Modeling recombination is difficult due to the large state space (recombination can potentially occur between any adjacent bases) and its non-Markovian nature.

## Data availability

The TGP data were downloaded from: https://www.internationalgenome.org/data-portal/data-collection/30x-grch38

The program MutAnce is available at: https://github.com/nage0178/MutAnce

Analysis scripts are available at: https://github.com/nage0178/mutInference_analysis

## Appendix

Let *Q* be the instantaneous rate matrix for a CTMC with 4 states where 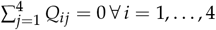. Define

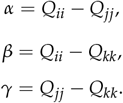

Let 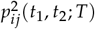 be the transition probability of going from state *i* to state *j* on a branch of length *T* via exactly two mutations that occur at times *t*_1_ and *t*_2_.

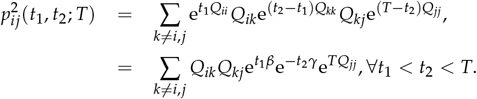

Similarly the probability a transition from state *i* to *j* on a branch of length *T* occurs via two mutations is given by,

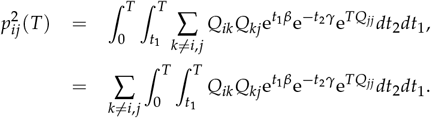

Here we assume it is legitimate to exchange the order of the sum and integrals. To derive a general expression for this equation, indicator functions are used,

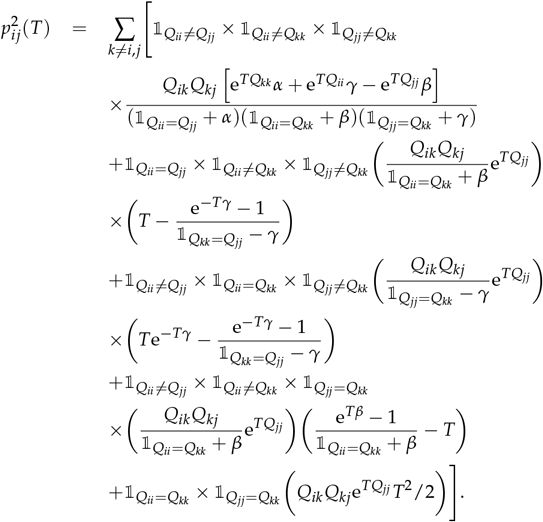

The joint probability density of the mutation time is then,

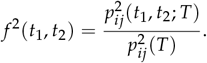

As we are only interested in observed mutations, if there are multiple mutations on a branch we find the marginal distribution for *t*_2_, as we can never observe the first mutation on a branch,

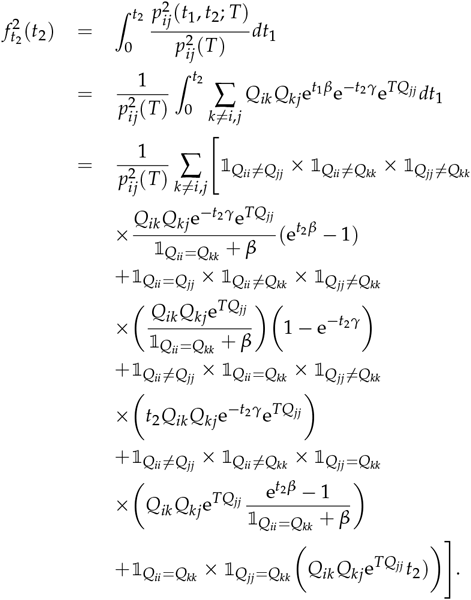

To simulate using the inverse transformation method we need the CDF,

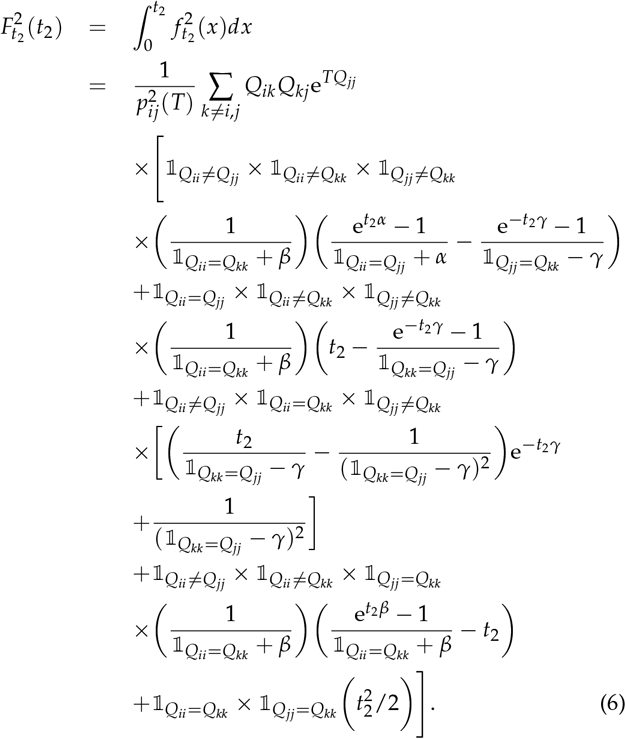

